# The Alzheimer’s disease neurodegenerative cascade reconstructed in human L2/3 excitatory neurons

**DOI:** 10.64898/2026.04.14.718430

**Authors:** Magdalena Zielonka, Anna Mallach, Bart De Strooper, Mark Fiers

## Abstract

Identifying the molecular cascade underlying neuronal degeneration in Alzheimer’s disease has been hampered by cellular heterogeneity and the limitations of donor-level classification. By integrating 851,682 cortical layer 2 and 3 excitatory neuron transcriptomes from 557 individuals across four independent snRNA-seq datasets of the human prefrontal cortex, we reconstruct neuronal degeneration as a continuous, stage-resolved transcriptional trajectory. Ordering neurons by collective pathological burden and clinical manifestation reveals that degeneration unfolds asynchronously within individual brains. Neurons in a single brain can simultaneously occupy early, intermediate, and late pathological states, a continuum entirely obscured by conventional approaches. The trajectory captures discrete transcriptional inflection points defining successive stages of vulnerability linked to development of neuropathology. Systematic analysis of the complete human kinome and phosphatome along this trajectory identifies a temporal hierarchy of phosphorylation dysregulation which collectively creates a permissive environment for tau pathology to escalate, defining stage-specific molecular nodes in the degenerative cascade.

## Introduction

Alzheimer’s disease (AD) is a complex progressive disorder characterized by amyloid-β (Aβ) plaques, neurofibrillary tau tangles (NFTs), gliosis, and ultimately neuronal dysfunction leading to cell death.^2–4^ Aβ pathology emerges early and is believed to exacerbate tau pathology. However, the molecular pathways linking these canonical hallmarks in vulnerable neurons remain incompletely understood.

Neuronal vulnerability in AD reflects the convergence of multiple pathogenic pathways, including disrupted glutamatergic signaling^5^, calcium dysregulation^6–8^, synaptic weakening, spine loss, oxidative stress^9,10^, mitochondrial impairment, ER stress, impaired autophagy, and tau hyperphosphorylation.^11–13^ These processes trigger micro- and astrogliosis, neuroinflammation, and regulated cell-death pathways such as necroptosis and senescence-like states^14–18^. Although each mechanism has been studied in depth, the temporal order and interactions between them in human tissue remain unresolved.

A central obstacle is the protracted prodromal phase of AD, during which molecular and cellular abnormalities accumulate slowly and asynchronously. Neurons within one individual therefore occupy diverse transcriptional and pathological states. Studies isolating NFT-positive and NFT-negative neurons show that only a minority of neurons exhibit overt pathology, each with distinct transcriptional programs^19^. Spatial transcriptomics further demonstrates that cellular responses vary according to proximity to Aβ plaques or tau tangles (Lu et al., in press)^20–22^. Yet conventional snRNA-sequencing typically aggregate neurons at the donor level, treating this biological heterogeneity as technical noise. This masks the continuum of neuronal states and obscures the underlying architecture of disease progression.

Among cortical neurons, layer 2/3 (L2/3) excitatory neurons consistently emerge as highly vulnerable in AD, showing early tau accumulation, transcriptional remodeling, and preferential depletion.^23–27^ Their extensive dendritic arbors, high spine density, and wide ranging excitatory connectivity enable higher-order cortical computations but impose substantial metabolic and homeostatic demands.^28–31^ These properties position L2/3 neurons as a critical entry point for reconstructing early neurodegenerative transitions in human AD.

To resolve the sequence of molecular events from healthy to diseased states, we integrated four high-quality snRNA-seq datasets (Lu et al., in press; Gazestani^24^; Gabitto^32^; Mathys^33^) yielding more than 850,000 L2/3 excitatory neuron transcriptomes. Using metacell aggregation we reduced inter- and intra-individual noise, achieved robust cross-cohort harmonization, and recovered reproducible pseudotime trajectories reflecting progressive neurodegeneration.^24,32,33^ Along this trajectory we identified discrete molecular inflection points associated with synaptic dysfunction, proteostatic stress, oxidative damage and survival pathway remodeling showing that the AD cascade can be reconstructed directly in human L2/3 neurons.

By resolving neuronal heterogeneity across over 850,000 L2/3 excitatory neurons, this framework reconstructs the sequence of transcriptional events that accompany human neuronal degeneration in AD at unprecedented resolution and offers a foundation for identifying therapeutic targets positioned at transitional stages of neuronal decline.

## Results

### Cross-dataset Integration of Layer 2/3 Excitatory Neurons

To reconstruct the transcriptional trajectory of neuronal degeneration at scale, we integrated snRNA-seq data from four independent studies of the human prefrontal cortex (PFC) (Gabitto, Gazestani, Mathys, and Lu (in press)).^24,32,33^. We selected the PFC to ensure regional consistency across datasets and because it is a region of progressive vulnerability in AD, where L2 and L3 excitatory neurons show pronounced synaptic loss and tau pathology. Three datasets (Gabitto, Mathys, and Lu) were generated from postmortem brain tissue, whereas Gazestani analyzed PFC biopsies from living individuals.

L2/3 neurons were initially defined using the original study annotations and were validated by reintegrating all excitatory neurons across datasets. This confirmed a robust, transcriptionally distinct, L2/3 cluster (Fig. S1A). We extracted this population for downstream analyses, excluding donors with insufficient cellular representation (Fig. S1B). The final integrated dataset comprised 851,682 L2/3 neuronal nuclei from 557 individuals, including cognitive healthy donors and donors spanning a broad range of AD stages (Figs. 1D-F) providing an unprecedented resource for investigating transcriptional changes associated with neuronal degeneration (Table S1).

**Figure 1.**
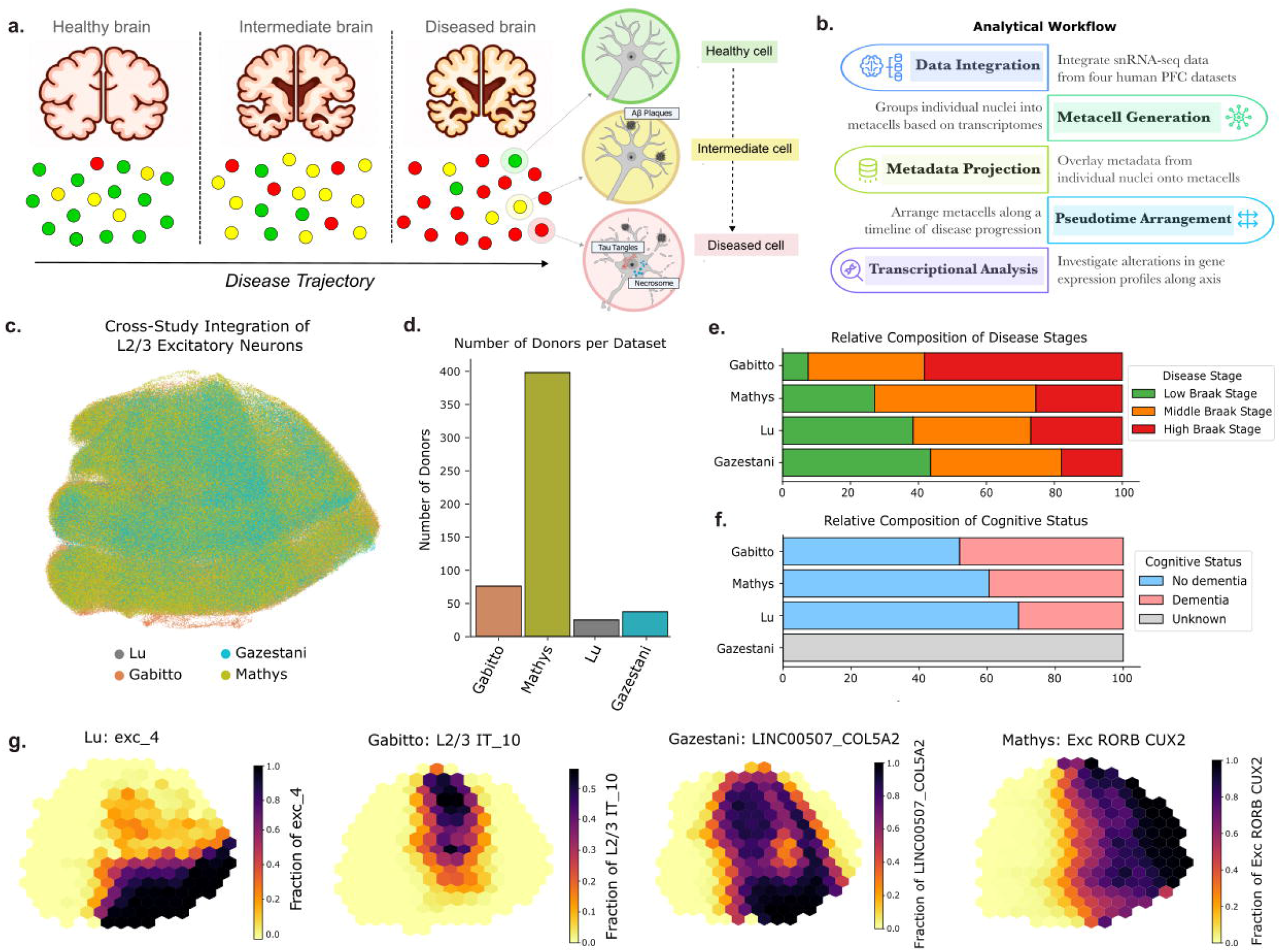
Integration of L2/3 excitatory neurons across datasets. **(a)** Schematic illustrating progression across the AD continuum, from a healthy brain with predominantly healthy neurons (green) to increasingly diseased conditions characterized by a heterogeneous mixture of healthy (green), intermediate (yellow), and degenerating (red) neurons. This progression reflects increasing amyloid and tau pathology and highlights the coexistence of neurons at different stages of the pathological cascade within the human brain. **(b)** Overview of analytical workflow. **(c)** Integrated UMAP embeddings of 851,682 single nuclei annotated as layer 2/3 excitatory (L2/3) neurons in their respective studies. Each point in the UMAP represents a single-nucleus RNA profile, colored by study. **(d)** Barplot showing distributions of donors from the four datasets. **(e-f)** Barplots showing relative distributions of donor disease stage (e) and cognitive status (f) across the four datasets (Low Braak stage: Braak stages 0-2 and Gazestani Control, Middle: Braak 3-4 and Gazestani Abeta, High: Braak 5-6 and Gazestani AbetaTau). **(g)** UMAP projections for each study, shown as hexbin density plots. Each hexbin represents a local aggregation of nuclei, with color intensity reflecting the proportion of nuclei annotated as the most vulnerable or responsive cell type in that study, highlighting differences in vulnerability designations across datasets.

We integrated the datasets (Fig. 1B) to correct for study-specific batch effects and to generate a unified UMAP embedding that captured the shared transcriptional structure of L2/3 neurons across all four datasets (Fig. 1C, Fig. S1C). Integration across cohorts highlighted notable differences in how neuronal vulnerability was defined across studies: subtypes designated as vulnerable in the original datasets did not consistently colocalize in the integrated transcriptional space, and disease status annotations showed similar variation across cohorts (Fig. 1G, Figs. S1D,E). Rather than reflecting intrinsic properties of the neurons, these differences largely arose from study-specific classification schemes, underscoring the value of a harmonized staging framework. We therefore unified donor-level disease status using Braak staging, the most consistently reported neuropathological criterion across studies, grouping donors into Low (<=2), Medium (3, 4), and High Braak (>=5) stages (Fig. 1E). Since Gazestani did not report Braak stages, we mapped donors according to the classification used in their study: control cases were designated “Low Braak,” Aβ cases to “Medium Braak,” and Aβ/Tau cases to “High Braak.” Donor cognitive status (dementia versus no dementia) was extracted from each study’s metadata, with the exception of Gazestani (Fig. 1F)

### Metacells reveal patterns of neuronal heterogeneity and disease association

We next applied the SEACell algorithm^34^ to aggregate transcriptionally similar nuclei into metacells, yielding 710 metacells for downstream analysis (Fig. 2A). Because each metacell contains nuclei drawn from multiple individuals and datasets (Fig. 2B, Figs. S2A,B), this approach integrates cross study metadata and captures gradual transcriptional shifts that span the disease continuum.

**Figure 2.**
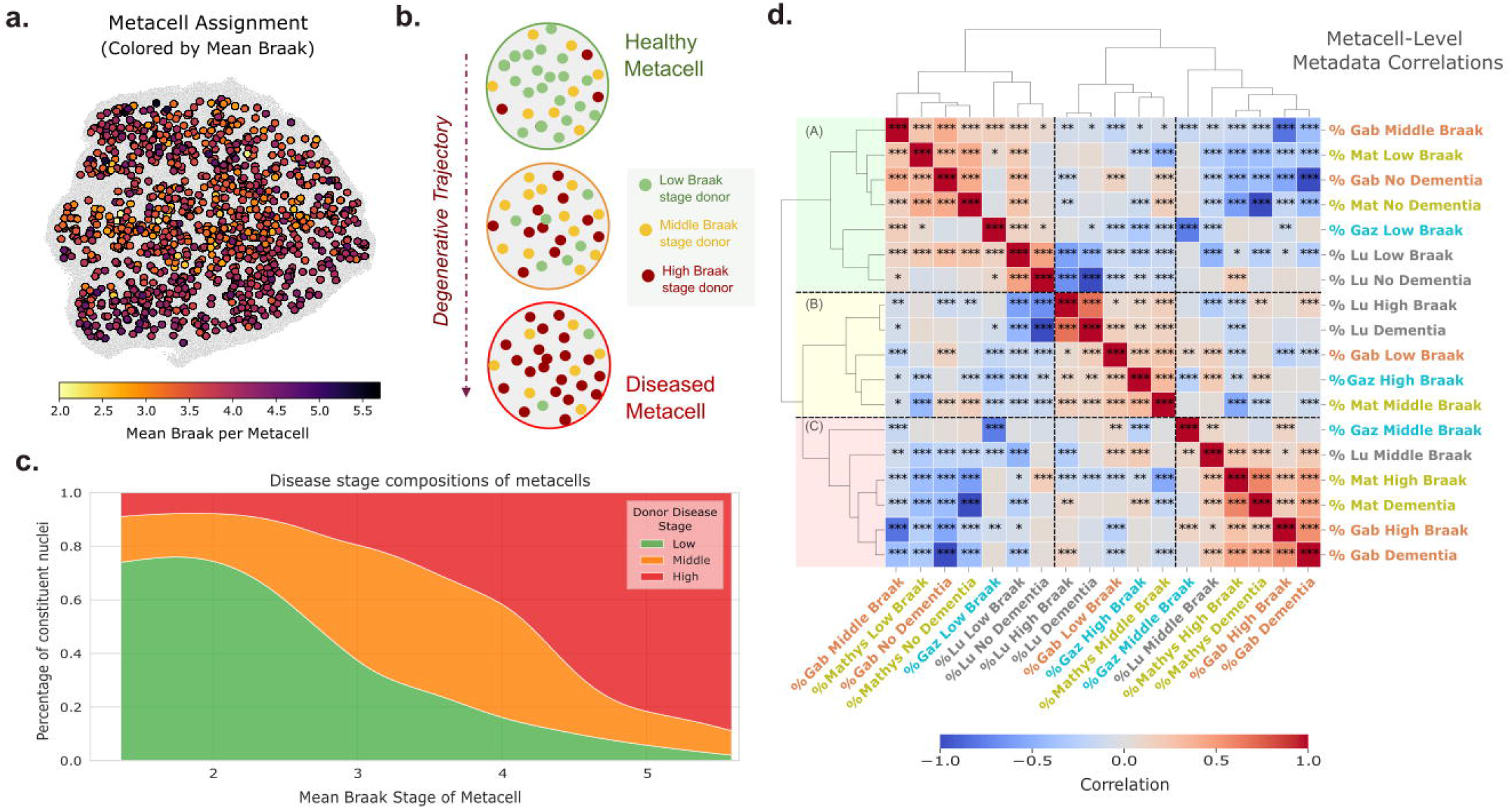
Transcriptomically defined metacells capture disease-stage heterogeneity. **(a)** UMAP of integrated single nuclei grouped into assigned metacells (n = 710), as generated by the SEACell algorithm. Small grey points represent individual nuclei, whereas larger, colored points represent aggregated metacells, colored by the mean Braak stage of the constituent nuclei. **(b)** Conceptual visualization of a healthy (top), intermediate (middle) and diseased (bottom) metacell; individual cells within each metacell are colored by donor disease stage. Metadata from individual cells can be aggregated to assign metacell-level scores. **(c)** Disease-stage composition of metacells. Metacells were arranged along the x-axis according to their mean Braak stage (based on donor-level Braak-stage of constituent nuclei). The y-axis denotes the fraction of cells within each SEACell originating from individuals at early, middle, or late stages. Generalized additive models were used to smooth stage proportions across the trajectory. **(e)** Heatmap of Spearman’s correlations between metacell composition scores, with scores defined as the proportion of constituent cells from each disease stage and dementia status category, stratified by study. Correlations between these scores highlight the relationships between donor disease stage and cognitive status across datasets within the metacells, demonstrating that disease-associated metadata compositions are concordant across independent cohorts and that metacells capture disease-relevant biological structure.

Aggregating metadata from constituent nuclei, including donor disease stage, cognitive status, and pathological burden, revealed that metacells with low or high mean Braak stage were strongly enriched for nuclei from healthy or diseased donors respectively, whereas intermediate metacells largely contained mixtures from multiple disease stages (Figs. 2A,C). Importantly, individual donors contributed nuclei to multiple metacell classes, indicating that these classes represent coexisting neuronal states within patients rather than mutually exclusive donor categories. This demonstrates that transcriptionally defined metacells capture neuronal states that progress within individuals, resolving a disease continuum that is obscured by donor-level classification.

To assess whether the association between metacell transcriptional state and disease status was reproducible across cohorts, we examined whether metacells enriched for disease-associated nuclei in one dataset showed concordant metadata compositions in the remaining datasets, correlating study-specific annotations across metacells in a clustered heatmap (Fig. 2D). Three major clusters emerged: two clusters (A and C) aligned strongly with healthy or diseased donor states across the cohorts, despite being defined solely by transcriptional similarity, whereas a third cluster (B) exhibited a mixed pattern, consistent with an intermediate or transitional state. The consistent correlation of disease-associated metadata compositions across independent cohorts demonstrates that disease-relevant biological structure is preserved within the metacell architecture, recapitulating structure in donor-level pathology while retaining intra-individual heterogeneity.

Together, these findings provide a harmonized cross-cohort framework in which L2/3 neurons can be analyzed along a continuous pathological trajectory, establishing the basis for reconstructing progressive transcriptional transitions during neuronal degeneration in AD (Fig. 2B).

### Ordering metacells along a metadata-informed disease continuum

Having established that metacells capture disease-relevant transcriptional structure, we next ordered them along a continuous trajectory that would directly link metacell transcriptional programs to progressive neuronal involvement in AD. We generated a metacell-level UMAP embedding based on the proportion of nuclei originating from donors at specific Braak stages or cognitive statuses, so as to incorporate both pathology burden and its clinical manifestation in the inferred trajectory. Applying diffusion pseudotime^35^ to the embedding revealed a single continuous trajectory, rooted in the metacell enriched for nuclei from cognitively healthy donors with low Braak stage (Fig. 3A). Along this trajectory, the proportion of nuclei derived from donors with high Braak stage and cognitive dementia increased steadily (Fig. 3B, Figs. S3A,B). Donor-level measurements of amyloid plaque burden and phosphorylated tau load (as measured in the Lu and Gabitto datasets) also rose monotonically along pseudotime (Fig. 3C), demonstrating that the transcriptionally inferred trajectory recapitulates the canonical pathological progression of AD, with amyloid accumulation preceding the emergence of tau pathology along the continuum.

**Figure 3.**
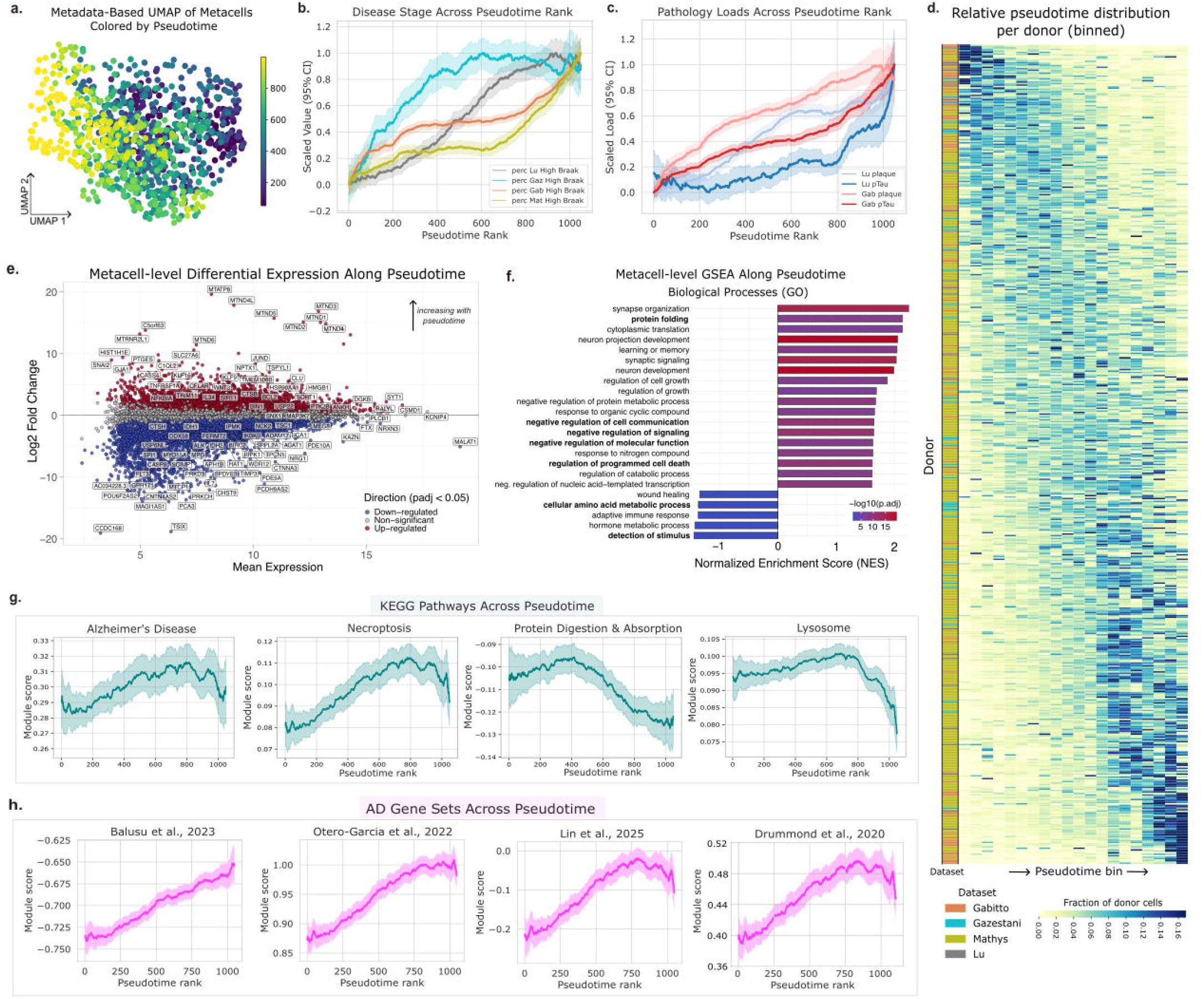
Pseudotemporal arrangement of metacells captures pathological progression and gene expression dynamics. **(a)** UMAP visualization of metacells, constructed from metadata features representing the percentage of constituent cells from each dataset assigned to specific disease stages or cognitive statuses. Diffusion pseudotime (DPT) analysis was performed on the embedding to identify continuous trajectories reflecting progressive shifts in metacell composition across disease progression. **(b)** Trajectories of metacell disease stage compositions along diffusion pseudotime. Metacell-level composition scores represent the proportion of nuclei in the metacell assigned to late disease stages. For each dataset the proportion of nuclei in the metacell assigned to late disease stages was standardized, smoothed using a centered rolling window, and plotted with shaded 95% confidence intervals of the mean. Smoothed trajectories were min–max scaled to facilitate comparison of relative progression patterns across datasets. **(c)** Trajectories of metacell pathology loads along diffusion pseudotime. Metacell-level values represent the mean donor-level pathology metrics derived from the Lu et al. and Gabitto et al. cohorts, averaged across nuclei within each metacell. For each dataset the pathology loads in each metacell was standardized, smoothed using a centered rolling window, and plotted with shaded 95% confidence intervals of the mean. Smoothed trajectories were min–max scaled to facilitate comparison of relative progression patterns across datasets. **(d)** Donor-level distribution of nuclei along the pseudotime trajectory. Metacells were grouped into 20 evenly spaced pseudotime bins based on pseudotime rank. For each donor (rows), the heatmap shows the fraction of that donor’s nuclei assigned to metacells within each pseudotime bin. Donors are ordered by median pseudotime, and the colored annotation bar indicates dataset membership. **(e)** MA plot of metacell-level differential expression across pseudotime. Each point represents a gene, with the x-axis indicating average expression and the y-axis representing the log fold-change along pseudotime rank. Genes above the y = 0 line are upregulated along pseudotime, while genes below are downregulated. **(f)** Gene set enrichment analyses (GSEAs) of genes differentially expressed across metacells with respect to pseudotime rank, showing most significantly enriched Gene Ontology Biological Processes (GO BPs). **(g)** Expression levels of selected KEGG pathways in metacells across pseudotime rank. Expression levels were calculated in metacells using scanpy’s *score_genes()* function. Values were locally smoothed using a rolling-window approach, with shaded regions representing 95% confidence intervals. **(h)** Expression levels of AD-associated gene sets, as identified in neuron-specific studies, in metacells across pseudotime rank. Expression levels were calculated in metacells using scanpy’s *score_genes()* function. Values were locally smoothed using a rolling-window approach, with shaded regions representing 95% confidence intervals.

Nuclei from individual donors were distributed continuously along the pseudotime axis, illustrating that degeneration proceeds at different rates across neurons within the same individual (Fig. 3D). Transcriptional heterogeneity was highest at intermediate pseudotime and converged toward both extremes, suggesting that neuronal states are most diverse during mid-stage disease transitions and become more uniform as neurons occupy either extremes of health and advanced pathology. Positioning nuclei along this continuous disease spectrum provides a more refined and biologically coherent view of progressive neuronal degeneration across donors, and implies that the window of greatest molecular diversity lies in the transitional stages before terminal degeneration is established.

### Transcriptional alterations along the pseudotime continuum

Differential expression analyses across metacells as a function of pseudotime identified a broad transcriptional shift from homeostatic to degenerative programs as neurons transitioned from healthy to diseased states (Fig. 3E, Table S2). Gene set enrichment analyses (GSEA) of pseudotime-associated genes revealed significant enrichment of Gene Ontology (GO) Biological Processes (BPs) and KEGG pathways linked to neuronal degeneration (Fig. 3F, Supplementary Fig. 3C, Table S3). Pathways related to regulation of cell death, neurodegeneration, protein folding, and negative regulation of cellular functions were among the most strongly upregulated, whereas metabolic pathways, stimulus detection, and immune response were downregulated.

Because neurodegenerative processes are unlikely to follow strictly monotonic dynamics, we sought to find more complex trajectories of key pathways along pseudotime by quantifying activity levels of pre-defined KEGG_2019_Human library pathways in each metacell and subsequently evaluating their behavior along the continuum (Fig. 3G, Fig. S3D). Some pathways, including Alzheimer’s Disease, necroptosis, endocytosis, and cellular senescence increased progressively along pseudotime, whereas others such as protein digestion and absorption showed a steadily decline. In contrast, pathways such as lysosome function and NF-κB signaling displayed non-monotonic patterns, demonstrating that certain disease mechanisms are not persistently activated but transiently engaged and subsequently suppressed along the degenerative cascade. Resolving neuronal states along a continuous trajectory rather than through conventional cross-sectional analyses makes these dynamics visible, revealing discrete stages of molecular vulnerability that would otherwise remain hidden within the degenerative cascade.

### Convergence of tau-associated signatures along the degenerative trajectory

Beyond tracking donor-level pathological burden, the trajectory captured the molecular hallmarks of tau pathology at the cellular level (Fig. 3H). Transcriptional and proteomic signatures (Table S4) associated with disease-relevant neuronal states—including NFT-bearing human neurons (genes differentially expressed in neurons with versus without neurofibrillary tangles)^19^, cortical pTau231 pathology (genes continuously varying with pTau231 burden across individuals)^36^, proteins physically interacting with PHF1-immunoreactive phosphorylated tau (as identified by affinity purification mass spectrometry)^36^, and human neurons xenografted into an AD mouse model (genes differentially expressed relative to control-grafted neurons)^37^–all increased progressively along the trajectory’ despite being derived from entirely independent experimental contexts. That gene expression profiles spanning post-mortem tissue, protein interactions, and cross-species models converge on a single transcriptional state suggests that the late-stage program identified here reflects a conserved molecular signature of tau-associated neuronal vulnerability rather than an artifact of any single experimental system.

### Temporal dynamics of gene expression programs reveal inflection points along the neuronal degeneration trajectory

To track coordinated transcriptional programs underlying this conserved trajectory of neuronal degeneration, we applied consensus non-negative matrix factorization (cNMF) ^38^, identifying 21 gene expression programs (GPs), that is, sets of co-regulated genes whose activity varies together across cells, rather than focusing on changes in individual transcripts (Figs. 4A,B, Supplementary Figs. 4A,B, Table S5). GSEAs of program-specific gene weights linked these programs to biologically interpretable processes, including mitochondrial function, synaptic signaling, protein homeostasis, lipid metabolism, axon guidance, and cell stress responses (Fig. 4E, Table S6).

**Figure 4.**
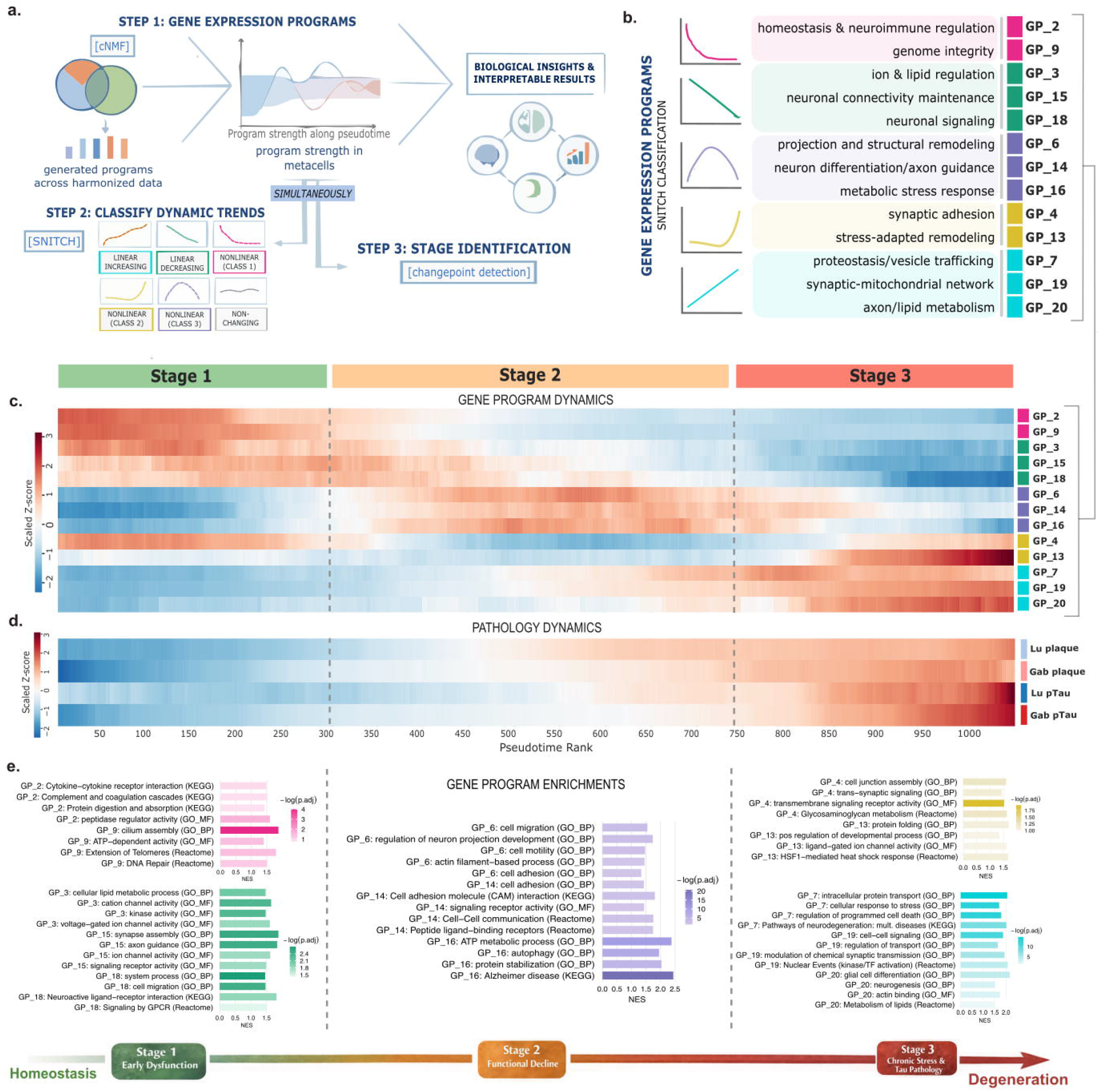
Temporal dynamics of metacells reveal coordinated transcriptional alterations along pseudotime. **(a)** Schematic of the workflow used for analyzing transcriptional dynamics of metacells along pseudotime. Co-expressing gene programs (GPs) were derived from the integrated dataset using cNMF, and their factor usage dynamics were tracked along pseudotime in metacells. Dynamic trends were classified with SNITCH, and multivariate change-point detection defined early, middle, and late stages, enabling interpretable insights into neuronal degeneration. **(b)** Classification of cNMF GPs according to SNITCH. Only GPs identified as having significant temporal shifts are visualized. GPs were named based on top pathway (GO and KEGG) enrichments (see panel (e)). **(c)** Heatmap showing sliding-window mean GP usage across metacells ordered by pseudotime, with colors representing scaled Z-scores of GP activity (red, high; blue, low). Pseudotime was partitioned into three stages (Stages 1–3) based on multivariate changepoint detection of coordinated shifts in mean gene program usage along the trajectory (**d)** Heatmap showing sliding-window mean pathology loads across metacells ordered by pseudotime, with colors representing scaled Z-scores of amyloid plaque and pTau measurements from the Lu et al. and Gabitto et al. datasets. **(e)** Pathway enrichments of cNMF GPs, based on gene set enrichment analysis of Gene Ontology terms as well as KEGG and Reactome pathways. GPs are grouped based on SNITCH classifications.

Mapping GP activity along the pseudotime trajectory, representing an inferred ordering of neurons along the degeneration continuum, revealed structured transcriptional changes underlying disease progression. Using the SNITCH algorithm^39^ we classified GP dynamics into four categories: linearly increasing (LI), linearly decreasing (LD), non-changing (NC), and non-linear (NL) (Figs. 4A-C, Table S7). Non-linear GPs were further grouped according to the direction and timing of their inflection points along pseudotime. Changepoint detection^40,41^ applied to GP activity trajectories revealed three transcriptional stages that mark progressive functional changes in neurons (Fig. 4C). These stages align closely with pathological progression: amyloid plaque accumulation predominates in stage 2, while tau pathology emerges more prominently in stage 3 (Fig. 4D, Fig. S4D).

The stage-specific dynamics of GP reflect coordinated functional transitions in neurons as degeneration progresses. Early along pseudotime, programs related to genome maintenance and synaptic regulation (GP_2, GP_9) decline rapidly, suggesting an early erosion of neuronal resilience. Membrane- and signaling programs (GP_3, GP_15, GP_18) decreased progressively, consistent with gradual disassembly of synaptic and structural functions. In contrast, stress-response and compensatory programs linked to proteostasis (GP_7), mitochondrial function (GP_19), and lipid metabolism/axon maintenance (GP_20), progressively increased, indicating adaptive responses to mounting cellular stress. Several GPs displayed transient or nonlinear dynamics (GP_6, GP_14, GP_16), suggesting early compensatory responses that ultimately fail to sustain neuronal homeostasis. Together, these analyses show that neuronal degeneration unfolds through coordinated transcriptional transitions marked by discrete inflection points in gene program activity, defining successive stages of vulnerability and stress adaptation along the AD trajectory.

### Stage specific kinase signaling accompanies progressive tau phosphorylation in vulnerable neurons

Closer inspection of the gene content of the two linearly increasing programs, GP_7 and GP_19, revealed that, embedded within their broader stress and mitochondrial signatures, both programs were highly driven by a notable subset of genes with mechanistic links to tau kinase regulation and phosphorylation state. GP_7 contained all five major cytoplasmic 14-3-3 isoforms (*YWHAG, YWHAH, YWHAB, YWHAE, YWHAZ*; phosphoserine/phosphothreonine-binding proteins found within NFTs in AD brain)^42^, co-expressed with *CDK5R1* (the activating subunit of the tau kinase CDK5), three calmodulin subunits (activators of the tau kinase CAMK2B), and *SET* and *ARPP19* (the endogenous inhibitors of PP2A, the principal neuronal tau phosphatase). GP_19 coupled this further to bioenergetic stress: *CAMK2B* and *GSK3A* were co-expressed within a program otherwise dominated by mitochondrially-encoded subunits of the oxidative phosphorylation machinery. Together, GP_7 and GP_19 implicated coordinated transcriptional remodeling of the tau phosphorylation machinery as a defining feature of the degenerative trajectory.

We therefore systematically classified pseudotime dynamics across the complete human kinase (kinome) and phosphatase (phosphatome) spectrum, using SNITCH, evaluating an unbiased set of kinases and phosphatases represented in the Brunello library^43^ and DEPOD database^43^, respectively (Fig. 5A, Table S8). Linearly increasing kinases included known tau kinases *CDK5* and *CAMK2B*, whose upstream regulators (*CDK5R1* and calmodulin genes, respectively) are encoded in GP_7 and GP_19, alongside *TTBK1, TAOK2, CSNK1E*, and *MAP2K7*. In contrast, kinases associated with neurotrophic and growth signaling, including *NTRK1/2, PIK3CB, MAP2K5, MAPK14, MTOR*, and *CDK7*, displayed declining trajectories. A third group, including *EIF2AK2, CAMK2A/G, AKT1, GSK3A, CSNK1G2*, and *PKN1*, displayed nonlinear trajectories whose activation onsets linearly increased until the Stage 2 to 3 boundary, at which point they plateaued.

**Figure 5.**
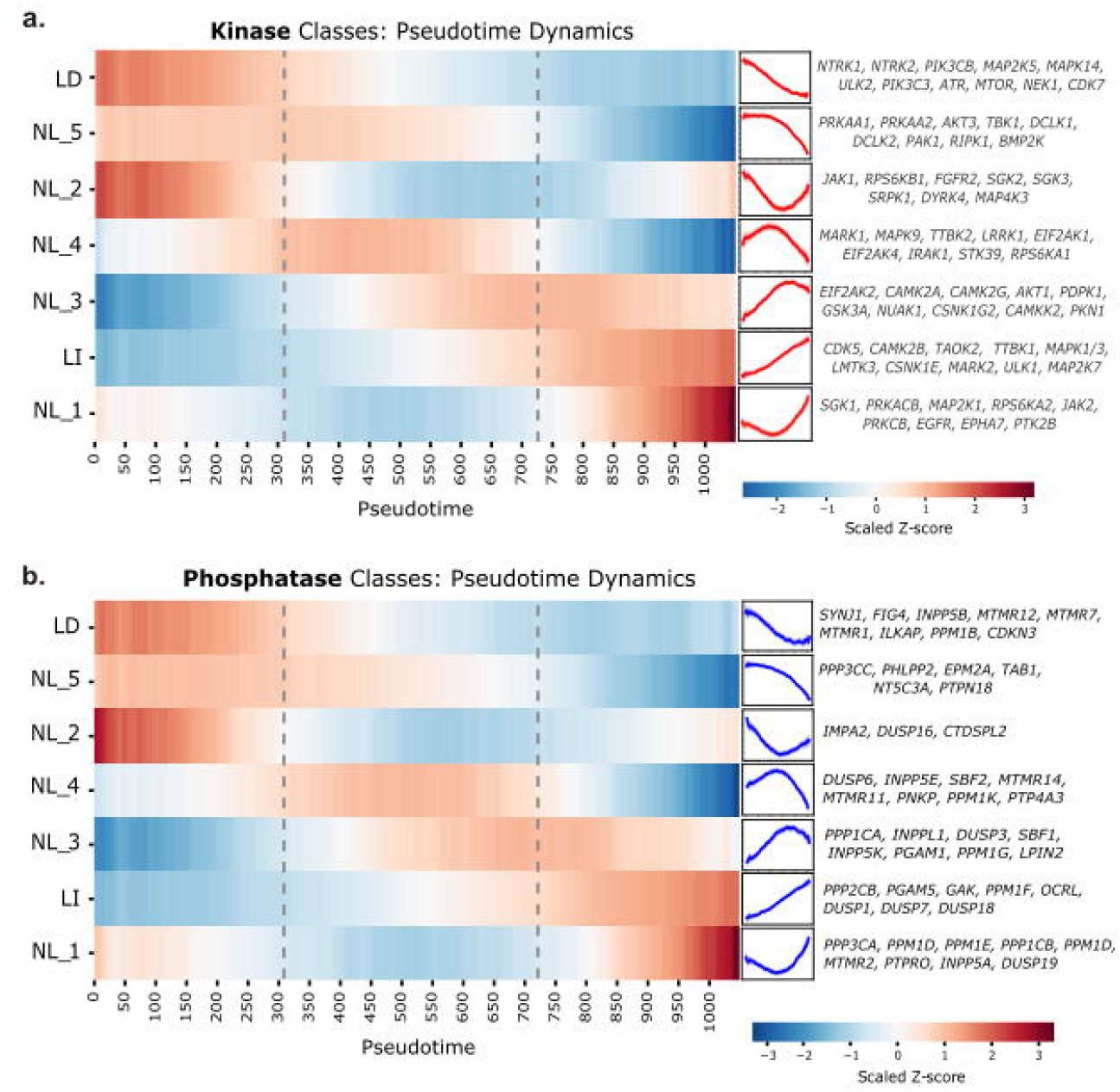
Temporal dynamics of kinase expression levels along pseudotime. **(a)** Heatmap (left) and lineplot (middle) of expression levels of each kinase class across metacells ordered by pseudotime. Classes were determined based on trajectory patterns using SNITCH. Class-level expression levels were calculated for metacells using Scanpy’s *score_genes()* function. Values were locally smoothed with a rolling-window approach. Representative kinases belonging to each class (right). **(b)** Heatmap (left) and lineplot (middle) of expression levels of each phosphatase class across metacells ordered by pseudotime. Classes were determined based on trajectory patterns using SNITCH. Class-level expression levels were calculated for metacells using SCANPY’s *score_genes()* function. Values were locally smoothed with a rolling-window approach. Representative kinases belonging to each class (right). Full list of categorized genes is available in Table S10.

Meanwhile, phosphoinositide phosphatases showed a coherent pattern where the majority of expressed family members declined linearly along pseudotime (*INPP5B, INPP1, MTMR12, FIG4, MTMR7, MTMR10, SYNJ1, MTMR1*) (Fig. S5A). Among linearly increasing phosphatases, the PP2A catalytic subunit *PPP2CB* rose progressively, while *PGAM5*, a mitochondria-localized phosphatase activated by bioenergetic stress, increased in parallel with the mitochondrial dysfunction encoded in GP_19. These analyses reveal that the phosphatome is dynamically reorganized along the degenerative trajectory in parallel with the kinome, with coordinated changes at both the family and individual gene level consistent with the progressive failure of endosomal, autophagic, and phosphorylation homeostasis in degenerating neurons.

## Discussion

Understanding L2/3 excitatory neuron degeneration in AD is complicated by cellular heterogeneity and inconsistent definitions of vulnerability. By integrating 851,682 L2/3 excitatory neuron transcriptomes across four independent cohorts and resolving them into a single pseudotemporal trajectory, we reconstruct the molecular cascade of neuronal degeneration in AD as a continuous, stage-resolved process. This framework reveals that degeneration does not unfold uniformly across individuals but asynchronously within them, as neurons in a single brain span early, intermediate, and late pathological states, a continuum that is entirely obscured by conventional donor-level classification. Aligning neurons according to their transcriptional state rather than donor identity disentangles intrinsic disease progression from inter-individual variability, revealing a coherent molecular cascade with discrete transcriptional inflection points that define successive stages of neuronal vulnerability.

When projected onto the shared pseudotemporal axis, divergent subtype annotations across datasets converge onto a consistent pattern: despite differences in clustering and nomenclature (e.g., L2/3 IT_10 in Gabitto versus exc4 in Lu), transcriptionally vulnerable neurons are progressively depleted, whereas disease-responsive states become enriched (Fig. S3E). This convergence not only validates the principal findings of each study but reveals that independently derived vulnerability annotations are actually concordant, complementary representations of a unified degenerative cascade. The divergence in subtype annotations across studies therefore reflects how study-specific clustering strategies attempt to divide what is, in reality, a continuous transcriptional spectrum into discrete classes. Unlike cluster-based staging, pseudotemporal ordering captures the full gradation of transcriptional change along the disease axis.

Within this unified trajectory, stage-resolved analysis delineates a temporally ordered sequence of molecular events. Early loss of programs supporting genomic maintenance, transcriptional control, and metabolic resilience (GP_9, GP_2) precedes excitability and synaptic dysfunction (GP_3, GP_15, GP_18), while an intermediate compensatory phase (GP_16) engages protein stabilization, autophagy, and metabolic support. In fact, this autophagic program is likely undermined by the simultaneous decline of *ULK2* and *PIK3C3*, suggesting that neurons attempt autophagic compensation transcriptionally but lack the kinase activity to sustain it.^44,45^ In parallel, GP_7 and GP_19 increase progressively from the earliest stages, reflecting a slow accumulation of proteostatic demand, chaperone activity, and mitochondrial strain that creates a continuously more permissive environment for tau hyperphosphorylation. By the terminal stage, neurons activate chromatin remodeling and mitochondrial quality-control pathways (GP_13), and induction of *HMGB1* signaling, a known damage associated molecular pattern, foreshadows cell death.^46^

That analysis reveals that the machinery of tau hyperphosphorylation is transcriptionally embedded within the neuronal stress response itself. The highest-weighted genes in GP_7 encode all five major cytoplasmic 14-3-3 isoforms (phosphoserine-binding proteins found within NFTs in AD) alongside the activating subunit of *CDK5*, the endogenous inhibitors of PP2A, and the calmodulins that activate *CAMK2B*.^42^ This supports the interpretation that GP_7 captures a transcriptional state with direct mechanistic relevance to tau pathology. Meanwhile, GP_19 deepens this further, as the co-expression of *CAMK2B* and *GSK3A* with mitochondrially-encoded OXPHOS subunits suggests that bioenergetic failure and tau kinase activation are coupled within the same program. The kinome screen extends this picture, identifying linearly increasing tau kinases including *TTBK1, TAOK2, CSNK1E*, and *CAMK2B*, while the concurrent rise of the PP2A catalytic subunit *PPP2CB* (alongside its endogenous inhibitors *SET* and *ARPP19* in GP_7) may reflect a compensatory transcriptional response to increasing tau phosphorylation. The early loss of metabolic resilience encoded in GP_9 and GP_2 is mirrored by the decline of neurotrophic kinases including *NTRK1, NTRK2*, and *MTOR*, while the coordinated decline of phosphoinositide phosphatases is consistent with the progressive endosomal trafficking dysfunction and autophagic impairment documented in AD neurons.^47,48^

The stage 2 to 3 boundary marks a discrete escalation superimposed on this continuous progression. *GSK3A* and *EIF2AK2* increase progressively and plateau at this boundary, suggesting that tau-directed kinase activation and integrated stress response engagement are sustained features of the most advanced neuronal states. Late-activated *PRKACB* encodes PKA, which directly phosphorylates tau and phosphorylates cAMP-regulated phosphoprotein (whose gene *ARPP19* is upregulated in GP_7), potentially perpetuating PP2A inhibition. The concurrent late activation of calcineurin (*PPP3CA*), which directly dephosphorylates tau, suggests a late counterregulatory response to offset the accumulated phosphorylation burden. Together, these findings define a temporal hierarchy in which the early coupling of tau kinase and phosphatase inhibitory machinery within GP_7 and GP_19, the progressive failure of neurotrophic, autophagic, and energy-sensing restraint, and late feedforward kinase activation collectively define the molecular logic of neuronal degeneration in AD.

This framework suggests stage-specific molecular nodes that may warrant further investigation. A prerequisite for identifying these nodes, however, is resolving the intra-individual heterogeneity that conventional donor-level analyses obscure. Even within the anatomically defined population of L2/3 excitatory neurons in the prefrontal cortex, neurons within a single brain simultaneously span early, intermediate, and late pathological states, indicating that disease stage is a property of individual neurons rather than of donors, with direct implications for therapeutic targeting and biomarker development. Future spatially resolved and perturbational studies will be critical to distinguish drivers from responses and to determine which of these nodes represent viable intervention points. More broadly, the trajectory framework developed here offers a generalizable approach for dissecting disease progression in other neurodegenerative conditions where cellular heterogeneity has obscured the underlying molecular sequence.

## Data availability

Datasets included in the manuscript are available in the following locations: (1) Gabitto et al.: through the <u>Open Data Registry</u> (https://registry.opendata.aws/allen-sea-ad-atlas/); (2) Gazestani et al.: through https://braincelldata.org/resource; (3) Mathys et al. through Synapse (https://www.synapse.org/#!Synapse:syn52293417). For reasons of ethics and privacy, snRNA-seq data from Lu et al. is deposited in the European Genome-phenome Archive (EGA) under study no. EGAS50000001692 with data accession no. EGAD50000002432. Requests for accessing raw sequencing reads must be submitted to EGA and will be reviewed by the VIB data access committee. For the purposes of this study, processed data from Lu et al. were kindly provided by the authors upon request.

An interactive viewer will be made publicly available upon publication, allowing users to explore metacell-level transcriptomic profiles across cell types and disease states, visualise gene set enrichment results, and query associated donor- and cell-level metadata. This resource is intended to facilitate exploration of the data presented in this study without requiring direct download of the full dataset.

## Code availability

The source code used in this study is available via GitHub at https://github.com/mmzielonka/Zielonka_NeuronalCascade_2026.

## Acknowledgments

This project received funding from the European Research Council (ERC) under the European Union’s Horizon 2020 Research and Innovation Program (grant agreement 101199326 XenoAD ERC-2024-AdG to BDS). This work was also supported by the Flanders Institute for Biotechnology (VIB vzw), a Methusalem grant from KU Leuven and the Flemish Government (METH/21/05 to B.D.S.), the Fonds voor Wetenschappelijk Onderzoek (G087523N to B.D.S.), the KU Leuven, the Queen Elisabeth Medical Foundation for Neurosciences (to B.D.S.), the Stichting Alzheimer Onderzoek (SAO-FRA 20240017 to M.F. and B.D.S.), the UK Dementia Research Institute (UK DRI-1004 to B.D.S.) and the Medical Research council (MR/Y014847/1 to B.D.S.), the Alzheimer’s Drug Discovery Foundation (to B.D.S.), the Alzheimer’s Association USA (to B.D.S), and the Edmond J. Safra Foundation through an Edmond and Lily Safra Fellowship (to A.M.).

We warmly thank Maria Livia Sassano and Dries T’Syen for scientific insights and discussions.

## Author contributions

M.Z., M.F., A.M., B.D.S. conceived the study. M.Z., M.F., A.M. developed the methodology. M.Z. analyzed the data. M.Z., M.F., A.M., B.D.S. drafted the manuscript. All authors reviewed and commented on the manuscript. M.Z. prepared the figures.

## Declaration of generative AI and AI-assisted technologies

During the preparation of this work, the authors used LLMs such as ChatGPT and Claude in order to support writing code for data analysis and to improve the readability and language of the manuscript. After using this tool or service, the authors reviewed and edited the content as needed and take full responsibility for the content of the publication.

## Supplemental information

**Document S1**. Figures S1-S5

**Table S1**. Donor-level metadata across four integrated datasets, related to Figures 1, 2, and S1.

**Table S2**. Metacell-level differential expression results across the pseudotime trajectory, related to Figure 3E.

**Table S3**. GO (Biological Processes) and KEGG enrichments identified by GSEA of metacell-level differential expression along pseudotime, related to Figures 3F and S3C.

**Table S4**. External AD-related gene set signatures, related to Figure 4H.

**Table S5**. Top 100 genes of each cNMF gene program, ranked by gene loading, related to Figure 4.

**Table S6**. GO biological processes, GO molecular functions, KEGG, Reactome, and WikiPathways enrichments of cNMF gene programs, related to Figure 4E.

**Table S7**. SNITCH classification of cNMF gene program trajectories based on average program usage in metacells along pseudotime, related to Figures 4 and S4.

**Table S8**. SNITCH classification of gene expression trajectories based on normalized expression in metacells along pseudotime, related to Figure 5.

## Methods

### Single-nucleus RNA-sequencing datasets

We analyzed four publicly available single-nucleus RNA-sequencing (snRNA-seq) datasets from postmortem human cortex (Gabitto et al., 2024^32^; Gazestani et al., 2023^24^; Mathys et al., 2023^33^), and an unpublished dataset from Nature Medicine (Lu et al., 2026, in press), for which processed data were kindly provided by the authors upon request. Analyses were restricted to excitatory neurons localized to cortical layers 2 and 3 (L2/3), and only individuals represented by at least 70 cells were retained to ensure robust modeling and reduce noise from sparsely sampled individuals.

## Quantification and statistical analysis

### Integration of datasets

All datasets were read into SCANPY.^50^ Genes expressed in fewer than 1% of cells per dataset were excluded. Count matrices were log-normalized using *sc*.*pp*.*normalize_total()* followed by *sc*.*pp*.*log1p()*, and highly variable genes were identified on the combined dataset using *sc*.*pp*.*highly_variable_genes()* (min_mean=0.0125, max_mean=3, min_disp=0.5, batch_key=‘dataset’). Batch effects were corrected using Harmony integration^51^ (*sc*.*external*.*pp*.*harmony_integrate()*) with “dataset” as the batch variable. Following initial integration, a small outlier cluster was removed, and the remaining cells were re-integrated to generate a harmonized L2/3 neuron dataset. For downstream analyses, individuals were assigned to Braak-stage groups: 0-2 as low, 3-4 as intermediate, and 5-6 as high. To enable direct comparison of neurons across cohorts while accounting for study-specific technical and annotation differences, we integrated the datasets (Fig. 1B) using Harmony to correct for the study as a batch effect. We then generated a joint UMAP^52^ embedding to capture the shared transcriptional structure across all four datasets (Fig. 1C, Fig. S1C).

### Visualization of cell densities on UMAP

To illustrate the spatial distribution and relative abundance of cells according to phenotype, cells were projected onto a two-dimensional UMAP embedding. Hexbin density plots were generated with a bin size corresponding to 20 hexagons per axis, in which each hexagonal bin represents the proportion of cells belonging to a given phenotype. A continuous color scale was used to indicate density, allowing for quantitative comparison of phenotype distributions across the embedding.

### Metacell construction

Metacells were generated from the integrated L2/3 neuron dataset using SEACells^34^, with one metacell constructed per ∼300 cells. Kernels were built on the Harmony-corrected PCA space, and archetypes were initialized using the first 10 eigenvectors and a waypoint proportion of 0.9. Metacells were iteratively refined until convergence (minimum 10, maximum 50 iterations; convergence threshold 1×10□□), and final SEACell assignments were summarized to generate metacell-level gene expression profiles. Metadata from constituent cells were aggregated to assign donor, phenotype, and other covariates at the metacell level for downstream analyses.

### Correlation analysis of metacell-level metadata

To examine relationships among cognitive, pathological, and study-specific variables, we computed pairwise Spearman correlations across metacell-level metadata. Spearman correlation coefficients and associated p-values were calculated using *scipy*.*stats*.*spearmanr()*, with missing values omitted. P-values were adjusted for multiple testing using the Benjamini–Hochberg false discovery rate procedure. Significant correlations were annotated according to conventional thresholds (p□<□0.05, *; p□< □0.01, **; p□< □0.001, ***). Correlation matrices were visualized as clustered heatmaps using *seaborn*.*clustermap()*.

### Pseudotime distributions across donors

To examine cell distributions along the inferred pseudotime trajectory, metacell-level pseudotime ranks were divided into 20 quantile-based bins. For each donor, the number of cells in each bin was normalized to the donor’s total cell count, yielding fractions per pseudotime bin. Donors were optionally ordered by median pseudotime. Braak-stage groups were indicated with a colored annotation strip alongside the heatmap, and an adjacent colorbar represented the fraction of cells per bin.

In parallel, the distribution of individual cells along pseudotime for each donor was visualized using Gaussian kernel density estimation (KDE, bandwidth□=□0.3). Densities were normalized per donor and stacked along the y-axis according to donor order, with donors colored by dataset of origin.

### Metacell feature space, dimensionality reduction, and pseudotime inference

To generate a robust metacell-level feature space, we used metacell-level metadata representing the proportion of cells in each cognitive status or Braak stage across datasets. Missing values were handled using an iterative PCA-based approach, followed by scaling and dimensionality reduction via PCA. The first 15 principal components were retained for downstream analyses, capturing ∼85% of the cumulative variance. These components were used to construct a neighborhood graph (*sc*.*pp*.*neighbors()*, 15 nearest neighbors) and generate a UMAP embedding (*sc*.*tl*.*umap()*) for visualization of metacell structure in low-dimensional space. Diffusion pseudotime (DPT)^35^ was then computed on this PCA/UMAP representation using *sc*.*tl*.*dpt()* to model cellular progression along the disease trajectory. The root cell was selected as the metacell with the lowest proportion of late-stage dementia cells and set as the root index.

### Rolling-window visualization of features along pseudotime

To visualize continuous changes in cellular features along the inferred pseudotime trajectory, nuclei were ordered by pseudotime rank and summarized using a rolling-window approach. For each feature—including sample metadata, pathology-associated measures, individual gene expression levels, or gene set activity scores—local averages were computed across neighboring nuclei using an adaptive window whose size varied along pseudotime, with smaller windows near the trajectory boundaries and a larger fixed window in the central region. Feature values were used in their original scale without transformation or normalization. At each pseudotime position, the rolling mean and associated 95% confidence interval were estimated using the standard error of the mean within the local window. Rolling trajectories were visualized as smooth curves along pseudotime, providing an intuitive representation of gradual, continuous changes across the disease spectrum.

### Metacell-level gene set scoring

To quantify the activity of gene sets and transcriptional signatures across the pseudotemporal trajectory, we computed gene set scores at the metacell level. Prior to scoring, raw counts for each nucleus were normalized to 50,000 total counts and log-transformed, then averaged across all nuclei within each metacell to generate metacell-level expression profiles. Normalization was applied at the single-nucleus level before metacell aggregation to ensure that differences in sequencing depth across datasets did not confound gene set activity scores. Gene set scores were subsequently computed on these metacell-level profiles using the *sc*.*tl*.*score_genes()* function.

### Consensus non-negative matrix factorization (cNMF)

To identify gene expression programs in □2/3 excitatory neuron metacells, RNA counts were preprocessed by selecting the top 4,000 highly variable genes, performing batch correction with Harmony (using dataset as the batch variable), and normalizing to TPM. Preprocessed data were input to cNMF^38^ for factorization with K values ranging from 16 to 25, 1000 iterations, and Frobenius reconstruction loss. The optimal number of factors (K□=21) was selected based on inspection of the factor stability and reconstruction error using the *k_selection_plot()* function. Following individual factorization runs, consensus gene programs were identified using cNMF’s clustering-based approach: program activity matrices from repeated runs were compared, and programs were grouped if they consistently co-clustered across runs. A density threshold of 0.08 was applied to define robust consensus programs, minimizing spurious or unstable components, and hierarchical clustering of programs was visualized to verify coherence. For each consensus program, every cell was massigned an activity score representing the relative contribution of that program to the cell’s transcriptome.

### Gene-level differential expression analysis

To identify genes whose expression changes along the disease-associated trajectory, we modeled expression counts for each gene as a function of metacell pseudotime. The gene expression profiles of all nuclei in each metacell were summed per SEACell. Gene expression from the resultant Seurat object was filtered to retain genes expressed in at least 4% of metacells. The fractions of cells in each metacell originating from each dataset were included as covariates. For each gene, a negative binomial generalized linear model (glmmTMB^53,54^) was fit with pseudotime rank as the primary predictor and metacell total counts as an offset.

### Gene Set Enrichment Analyses (GSEA)

To identify biological processes and pathways enriched along the pseudotime trajectory, genes were first ranked by their standardized log fold change. The ranked list was used as input for gene set enrichment analyses (GSEAs)^55^ using the clusterProfiler^56^ R package, assessing enrichment of Gene Ontology biological processes (GO BPs)^57,58^ and KEGG^59^ pathways. Resultant p-values were corrected for multiple testing using a Benjamini-Hochberg correction. To reduce redundancy among enriched GO terms, semantic similarity-based clustering was performed using the *simplify()* function in clusterProfiler, applying the Wang similarity measure with a cutoff of 0.6 and retaining representative terms based on the lowest adjusted p-value.

### Trajectory classification using SNITCH

To systematically classify the pseudotime dynamics of gene expression programs and individual genes, we applied SNITCH^39^, a computational framework that detects linear and nonlinear trajectory patterns along a continuous axis. SNITCH fits both linear models and generalized additive models (GAMs) to each feature’s trajectory, uses the Bayesian Information Criterion to select the best-fitting model, and assigns each feature to one of the following trajectory classes: linearly increasing (LI), linearly decreasing (LD), nonlinear (NL), variance increasing (VI), or non-correlated (NC). Nonlinear trajectories are further subclustered using unsupervised clustering of functional principal component analysis (FPCA) representations to identify distinct nonlinear patterns, and all classifications are corrected for multiple testing using the Benjamini-Hochberg false discovery rate procedure. SNITCH was applied to two feature sets along the metacell pseudotime trajectory: (1) to the 21 cNMF gene program activity scores computed at the metacell level and (2) to was applied to individual gene expression values across (normalized as described in metacells. Prior to SNITCH classification, gene expression values were normalized as described in the “Metacell-level gene set scoring” section.

### Multivariate changepoint detection and stage delineation

Pseudotime-ordered metacell activity scores for each of the 21 cNMF gene programs were summarized using a sliding window with a dynamic window size ramping from 200 to 400 cells, yielding smoothed mean trajectories with 95% confidence intervals. Each trajectory was baseline-shifted to zero and scaled by its maximum absolute value to enable cross-program comparison. The smoothed activity scores of all 21 programs were then stacked into a multivariate signal matrix, z-score normalized, and submitted to multivariate changepoint detection using the PELT algorithm^60,61^ with a radial basis function cost model as implemented in the ruptures package. The penalty parameter was set to 100 and detected breakpoints were mapped back to pseudotime indices to define discrete trajectory stages.

