## Supplementary material for "The Alzheimer’s disease neurodegenerative cascade reconstructed in human L2/3 excitatory neurons": Document S1

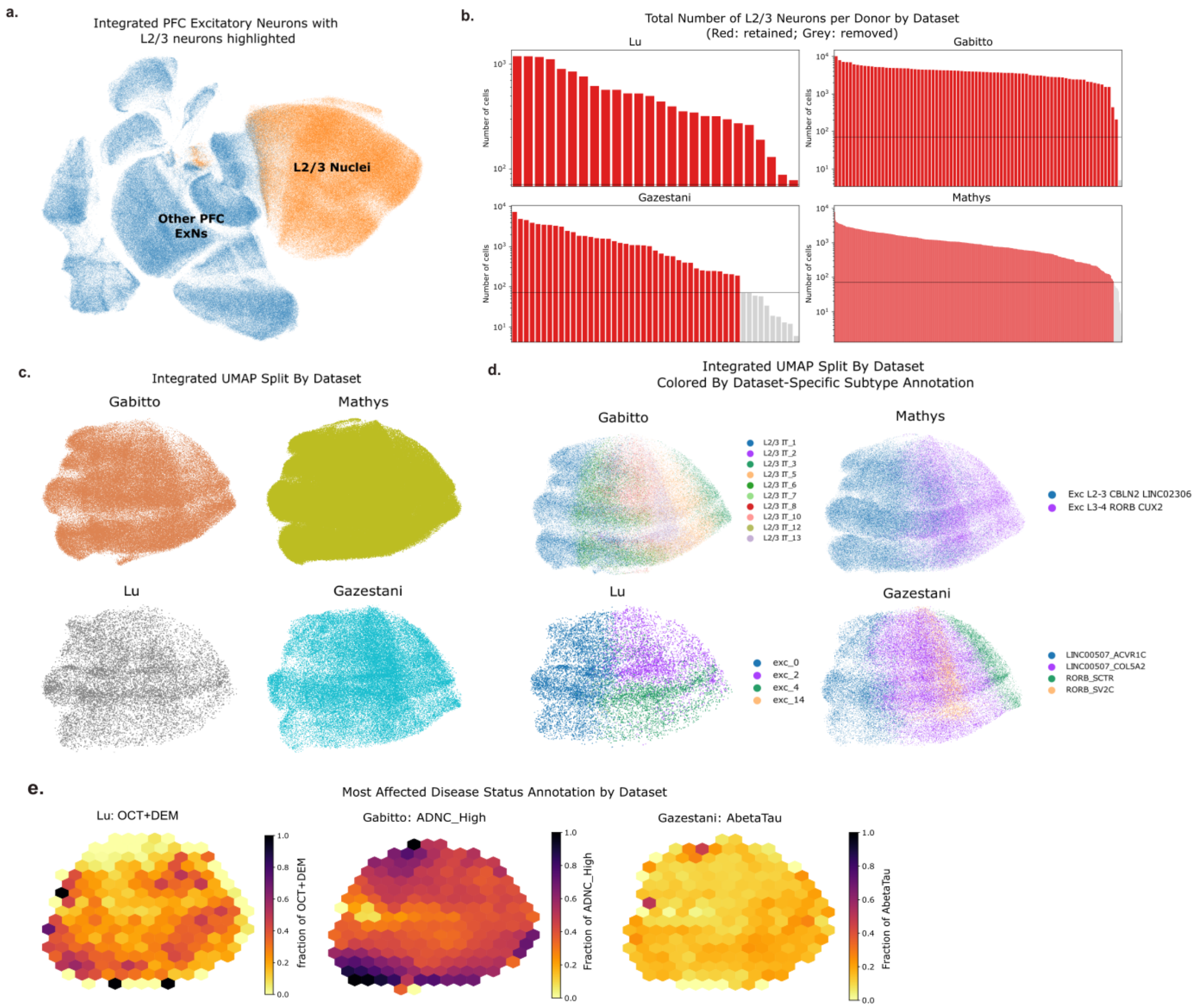


**Supplementary Figure 1. Integration and characterization of layer 2/3 excitatory neurons across datasets**

**(a)** Integrated UMAP embeddings of all excitatory neurons extracted from the four datasets, with single nuclei annotated as layer 2/3 excitatory neurons in their respective studies highlighted in orange. Each point in the UMAP represents a single-nucleus RNA profile. Note that, given the representative nature of this figure and the computational demands of full dataset integration, only a subset of 54 individuals was included from the Mathys et. al dataset. **(b)** Number of L2/3 excitatory neurons obtained per individual, stratified by dataset. Only donors with at least 70 nuclei were included in the downstream analysis to ensure adequate representation of donors. **(c)** Integrated UMAP embeddings of L2/3 neurons, grouped by dataset. **(d)** Integrated UMAP embeddings of L2/3 neurons, grouped by dataset and colored by original cell subtype annotation, as assigned in the original publication. **(e)** UMAP projections for each study, shown as hexbin density plots. Each hexbin represents a local aggregation of nuclei, with color intensity reflecting the proportion derived from donors classified as most affected in that study.

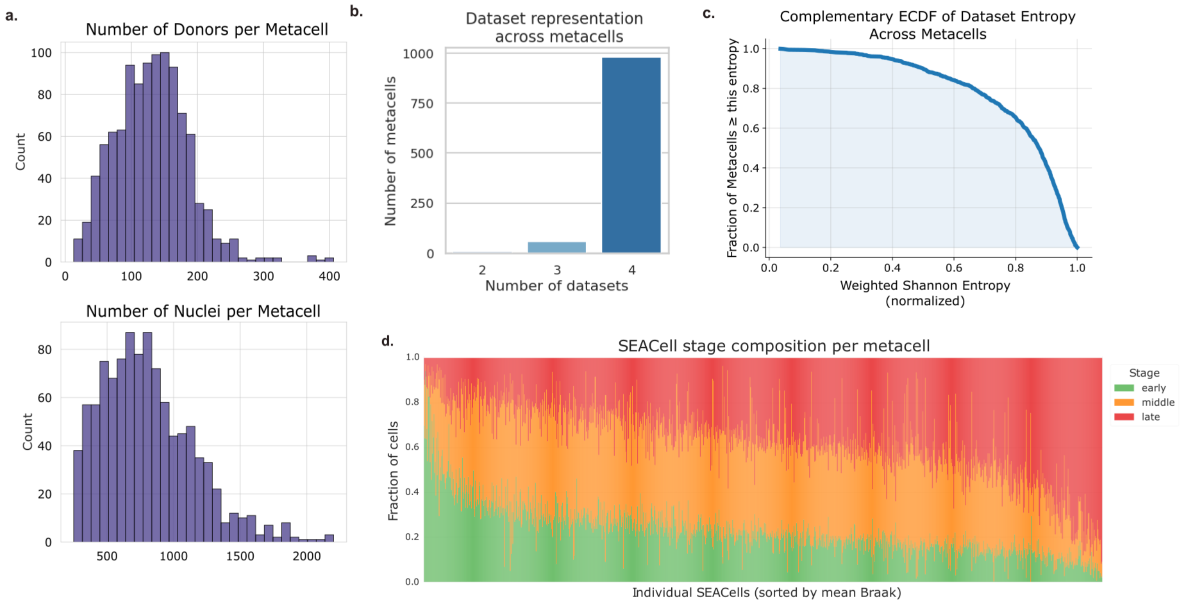


**Supplementary Figure 2. Diversity and disease-stage representation across metacells**
**(a)** Distribution of number of nuclei contained in each metacell (top) and distribution of number of unique donors represented in each metacell (bottom). **(b)** Barplot showing number of datasets represented across each of the metacells. **(c)** Complementary ECDF of metacell dataset entropy. Normalized Shannon entropy across metacells, reflecting the diversity of dataset contributions. The complementary ECDF shows the fraction of metacells with entropy ≥ a given value. **(d)** Metacell disease-stage composition. Stacked bars show the fraction of nuclei from early (green), middle (orange), and late (red) Braak stages within each metacell, ordered by mean Braak stage.

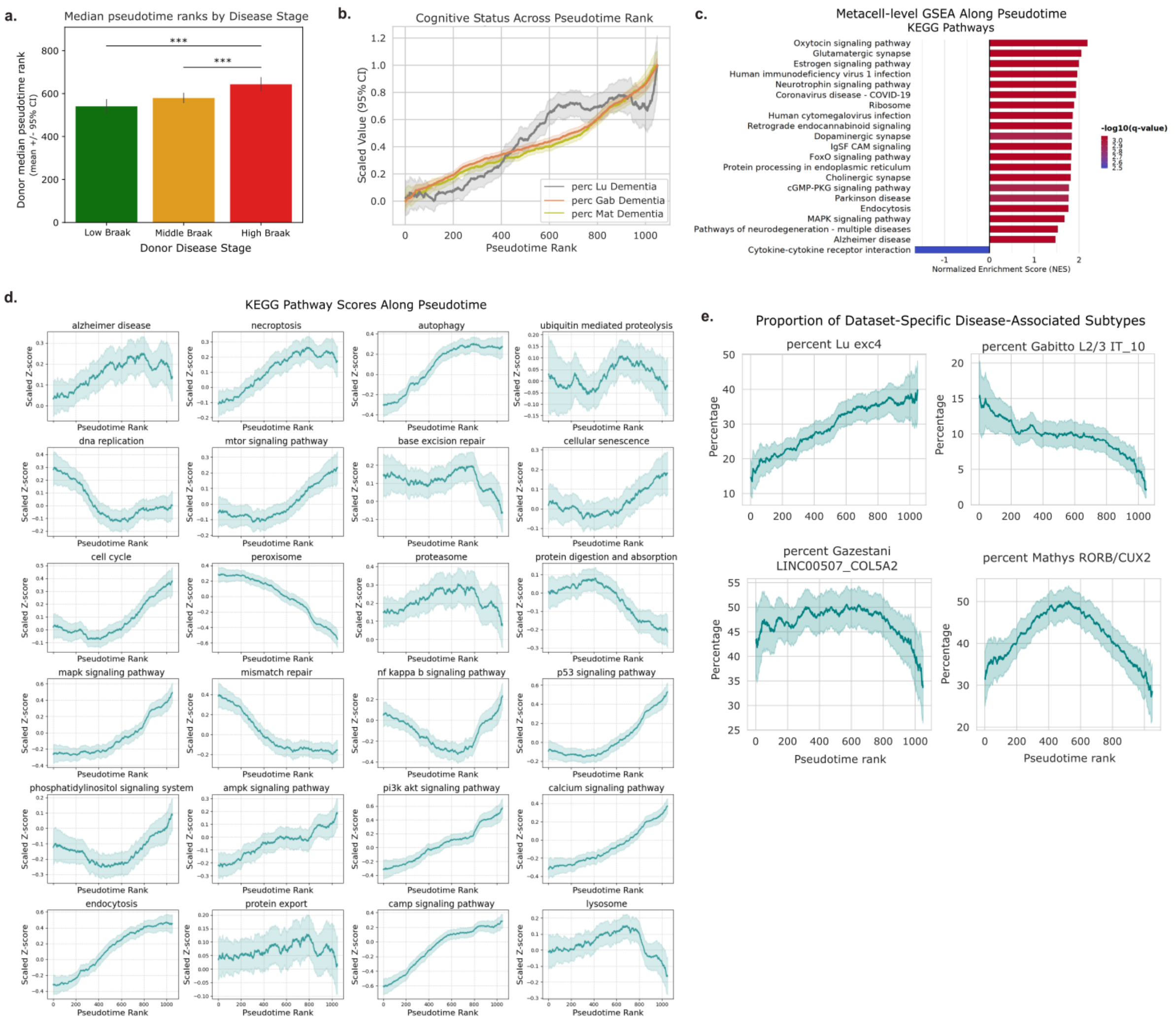


**Supplementary Figure 3. Pseudotemporal dynamics of metacell composition, cognitive status, and KEGG pathway activity**

**(a)** Bar plot showing median pseudotime rank of metacells grouped by disease stage (green, orange, red). Bars indicate group means with 95% confidence intervals. Differences between stages were tested using one-way ANOVA, followed by Tukey's HSD post hoc test ($\alpha = 0.05$). Significant pairwise comparisons are indicated by asterisks. **(b)** Trajectories of metacell "cognitive status" compositions along diffusion pseudotime. Metacell-level composition scores represent the proportion of nuclei in the metacell originating from donors classified as having dementia. For each dataset the proportion of nuclei in the metacell assigned to dementia was standardized, smoothed using a centered rolling window, and plotted with shaded 95% confidence intervals of the mean. Smoothed trajectories were min–max scaled to facilitate comparison of relative progression patterns across datasets. **(c)** Gene set enrichment analyses (GSEAs) of genes differentially expressed across metacells with respect to pseudotime rank, showing most significantly enriched KEGG pathways. **(d)** Expression levels of selected KEGG pathways in metacells across pseudotime rank. **(e)** Trajectories of metacell vulnerable/responsive cell type composition along pseudotime. Metacell-level composition scores represent the proportion of nuclei annotated as the most disease-vulnerable or -responsive excitatory neuron subtype, as defined independently in each source study. For each dataset, proportions were standardized, smoothed using a centered rolling window, and plotted with shaded 95% confidence intervals of the mean.

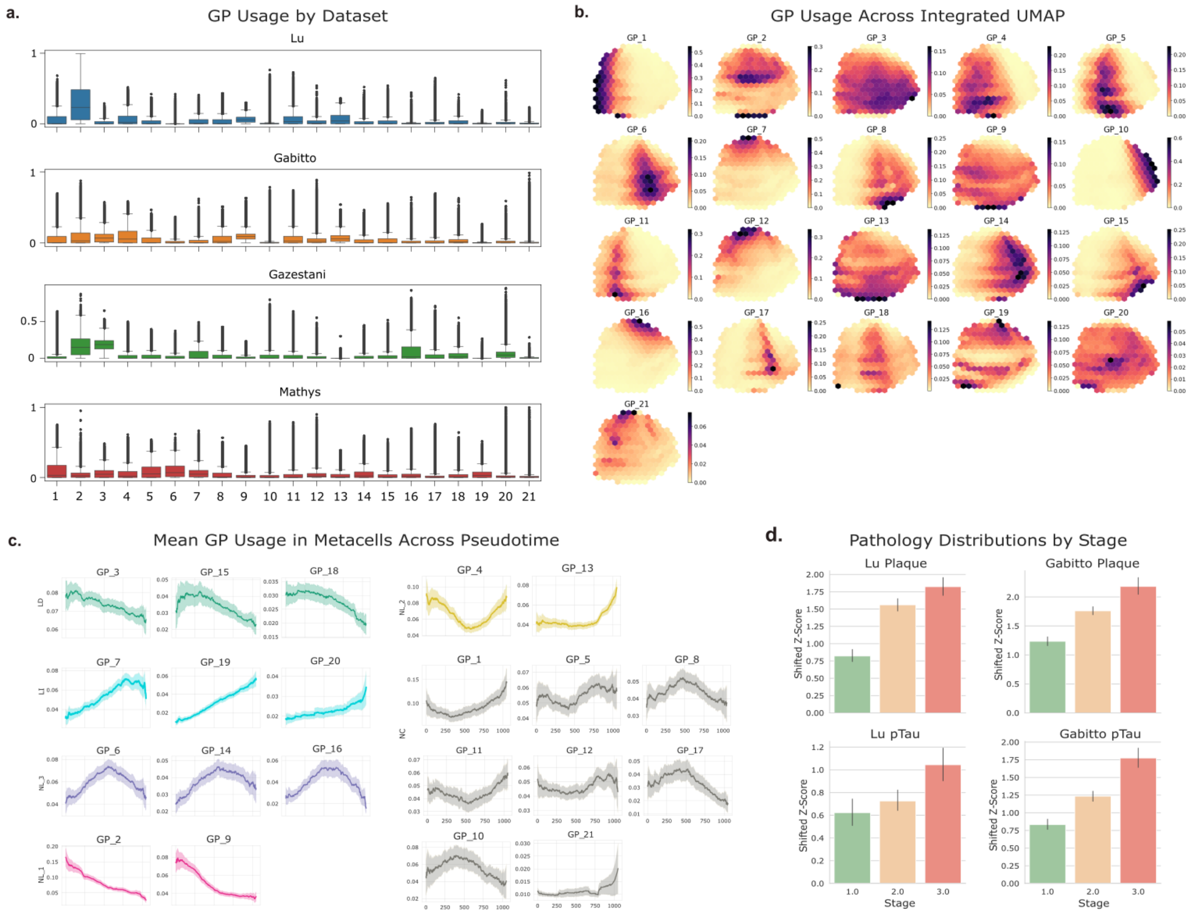


**Supplementary Figure 4. Pseudotemporal patterns of cNMF gene program usage and pathology in metacells**

**(a)** Distribution of per-cell usage scores for each of the 21 cNMF gene programs (GPs), split by dataset. **(b)** Hexbin density plots of cNMF GP usage projected onto the integrated UMAP, with color intensity reflecting usage score. **(c)** Trajectories of metacell GP usage scores along diffusion pseudotime. GPs are categorized according to their SNITCH classification. **(d)** Bar plots showing mean pathology values for amyloid plaque and pTau measurements from the Lu et al. and Gabitto et al. datasets, grouped by pseudotime stage (as determined in Fig. 4c). Bars represent the scaled mean value per stage, with 95% confidence intervals indicated by error bars.
